# Investigation of the global transportation of *Culicoides* biting midges, vectors of livestock and equid arboviruses, from flower-packing plants in Kenya

**DOI:** 10.1101/2025.04.01.646514

**Authors:** J.E. Stokes, K. Labuschagne, E.M. Fèvre, M. Baylis

## Abstract

**Background:** In recent decades there has been a huge increase in the export of cut flowers from countries in Africa and elsewhere to European flower markets, with the vast majority first entering the Netherlands for local use or for export. Coincidentally, two significant livestock disease outbreaks caused by viruses associated with Africa or other tropical regions, were first detected in the Netherlands (bluetongue virus, BTV, 2006) and in western Germany about 200 km from the Netherlands border (Schmallenberg virus, SBV, 2011). This study aimed to determine whether *Culicoides* biting midges, the vectors of BTV and SBV, are present within flower-packaging plants in East Africa, and therefore whether *Culicoides* could be unknowingly exported during the shipping of cut flowers.

**Methods:** Field sampling was undertaken at a flower-packaging facility in Kenya, East Africa. The facility undertook all stages of cut flower production from maintaining rootstock through to packaging and shipping to an airport for international export. Trapping was undertaken at each stage of production (rootstock, propagation, inside growing greenhouses, in the packing-house, inside cold-storage rooms, during transportation) using CDC Light Emitting Diode (LED) light traps. Hand-held aspirators were used to obtain individual insects directly from flowers and around composting sites, while emergence traps studied insect emergence from compost, leaf-litter and flowers discarded at quality control checkpoints.

**Results:** A maximum nightly catch of 269 *Culicoides* were identified on a half-acre smallholding, containing 15 ruminants and 40 birds, located 20 m from the nearest greenhouse. Greatest numbers of *Culicoides* were trapped at a pond (*n =* 23) and leaf-litter compost site (*n =* 19) within the curtilage of the flower-packaging plant. Of the seven greenhouses sampled, three had *Culicoides* trapped overnight (mean = 4, range: 1-9), and no *Culicoides* were trapped in the propagation units. No *Culicoides* were trapped in the pack house, cold-store, or during transportation of the flowers to the airport for shipment. No *Culicoides* emerged from emergence traps or were trapped when aspirating directly from flowers.

**Conclusions:** This is the first study to investigate whether *Culicoides* are present within flower packaging plants in Africa. The results highlight that although present in small numbers both outside and within greenhouses, the presence of *Culicoides* declined with each stage of production. Therefore, the risk of exporting *Culicoides* with packaged cut flowers is non-zero but likely very small.

## Background

Biting midges of the genus *Culicoides* (Diptera: Ceratopogonidae) transmit several important viruses of ruminants, such as bluetongue virus (BTV), Epizootic Haemorrhagic Disease virus and Schmallenberg virus (SBV) as well as African horse sickness virus [1]. Prior to 1998, BTV was considered exotic to Europe, with subsequent incursions of several BTV serotypes into southern Europe between 1998 and 2006, limited in distribution to the Mediterranean Basin where the main Afro-Asiatic vector, *Culicoides imicola*, was present [2]. However, in August 2006, a BTV serotype 8 (BTV-8) outbreak was detected in Limburg Province in southern Netherlands [3], from where it quickly spread throughout Europe, demonstrating the potential for European Palearctic *Culicoides* species to sustain and propagate outbreaks. The BTV strain occurred 900 km further north than the northern-most limit of previous European BTV incursions and entirely bypassed southern Europe. Full genome sequencing of the virus indicated a sub-Saharan origin [4].

In 2011, a novel *Culicoides*-borne disease was detected near the eponymous town of Schmallenberg in western Germany, only ∼200 km from the original point of detection of BTV-8 five years earlier [5]. Schmallenberg disease causes mild clinical signs in adult cattle and sheep but can cause congenital malformations and stillbirths in both species if infection coincides with the vulnerable stages of gestation [6]. The causative agent (Schmallenberg virus, SBV) was a novel Simbu serogroup virus from the genus *Orthobunyavirus*; with sequence analysis suggesting SBV is a reassortant of Shamonda and Sathuperi viruses [7]. Although some orthobunyaviruses had previously been reported in Europe, viruses from the Simbu serogroup had not previously been isolated in the region but are widespread in other parts of the world [8].

There have recently been further incursions of BTV into the Netherlands. In September 2023 BTV-3 was detected in sheep in central Netherlands, about 30 km southeast of Amsterdam and a similar distance from Schipol, its airport [9]. In October 2024 BTV-12 was detected in a sheep, close to where BTV-3 had first been detected [10]. The origins of these viruses are unknown at present.

The emergence of these viruses highlights the need for a greater understanding of potential incursion routes of *Culicoides*-borne viruses into northern Europe. An investigation of possible entry routes of BTV-8 into northern Europe in 2006 considered the legal or illegal import of infected animals; the import of biological materials potentially containing the virus (vaccines, serum, semen); or the import of infected *Culicoides*, either on the wind, in aircraft or packed up with plants [11]. The study failed to identify the route conclusively but suggested that the potential import of infected *Culicoides* should be further studied.

Here we investigate the potential for *Culicoides* to be imported to northern Europe with cut flowers. Flowers exported from Africa to Europe are grown in greenhouses, packed at night under bright lights [11], wrapped in polythene, chilled and transported by air. It is theoretically possible that *Culicoides* could be packed up with the flowers, survive the journey and be released after arrival. Routine monitoring of pests in exported plants is not undertaken either by farms or by exporters as it would disrupt the just-in-time supply chain for fresh products such as flowers, and export controls are largely risk-based.

The international trade in cut flowers has grown dramatically in recent years, with an export value of $10 Bn in 2022 [12]. The flowers are primarily sourced from Latin America and Africa and supply markets in North America and Europe. In Europe, the Netherlands is a hugely significant hub with, in 2014, about 60% of all imports of cut flowers to the European Union entering the Netherlands first, before either local purchase or export to other countries [13].

For example, in 2017 the Netherlands imported nearly 273 thousand tons of cut flowers. Pests are regularly found in consignments: an investigation of cut flowers imported into Australia found about forty percent of consignments from Kenya harboured pests [14]. An analysis of 1,538 interceptions into the UK and the Netherlands in 2017 identified 459 different pest types, about 90% insects and ∼6% arachnids [12]. The vast majority were plant pests, such as white fly, noctuid moths, leaf miners, thrips and aphids. However, it is noteworthy that in the Netherlands both Culicidae (the family of mosquitoes) and Nematocera (a suborder of Diptera, that includes mosquitoes and midges) were listed amongst identified pests.

This study aims to test the hypotheses that female *Culicoides* 1. can be found within flower-packaging plants in Kenya supplying goods to global flower markets; 2. are found during each stage of the flower production and packaging process; and 3. are found during transportation from flower-packaging plants to the airport.

## Methods

### Field Site

Field sampling was undertaken during October 2014 at a flower-growing and packaging facility in Kenya, East Africa, at 1,500 m altitude. The precise location of the farm cannot be disclosed, to protect commercial interests. The farm undertook all stages of flower production from maintaining rootstock and outdoor crops, to propagation of young stock and growing on of established plants, to packaging flowers to individual customer requirements and exportation of packaged products using cold-storage methods. Flowers which did not meet quality control expectations were composted on site. Although more than one variety of flower was grown, the farm specialised in the exportation of roses into the European market, to both auction houses as well as directly to supermarkets and wholesalers. More than twenty greenhouses were used to grow roses on site, varying in size between a 0.3 hectare ‘show’ greenhouse, used to show customers the full variety of roses grown on site, up to 2.7 hectare greenhouses for growing-on the most popular rose varieties. The total size of the greenhouses was 60 hectares (0.6 km^2^)

### Field Sampling

Trapping was undertaken over eight nights using Centers for Disease Control and Prevention (CDC) Light Emitting Diode (LED) light traps connected to a 6-volt battery. Sixteen sites were selected to represent each stage of production (rootstock, propagation, inside growing greenhouses, in the packaging-house, inside cold-storage rooms, during transportation). Each site was sampled for 2 nights (with 4 sites sampled each night over the 8-night period). Hand-held aspirators were used to obtain individual insects directly from flowers and around composting sites, while emergence traps were used to capture any freshly emerging insects from compost, leaf-litter and flowers discarded at quality control checkpoints. Details of the type of production system, as well as the use of any insecticides, and biological and mechanical control methods used within each greenhouse, were also recorded.

### Smallholdings

Additionally, two smallholdings (hereafter referred to as SH1 & SH2) that directly neighboured the facility had CDC traps present for 3 nights each. SH1 was located 270 m from the nearest greenhouse at the flower farm and contained 10 dairy cattle, a small number of dairy goats and 30 rabbits on 2.5 hectares. The majority of SH1 was used for field crops supplied by drip-irrigation and vermi-culture compost. SH2 was 20 m from the nearest greenhouse and contained 15 ruminants and 40 birds, on a 0.38 hectare site

### Specimen Identification

*Culicoides* were separated from other insects according to their wing characteristics using a stereomicroscope and stored in 70% ethanol. *Culicoides* were identified to species level based on their wing marking; other insects were mostly identified to the level of family or order.

## Results

### Insects Trapped

The 38 trap catches produced a total of 445 *Culicoides* of 13 species, and 2,617 other insects of 26 genera over the eight nights of trapping. The single largest nightly catch of *Culicoides* was 269 at SH1 (140 *C. imicola* (3 male), 91 *C. bolitinos*, 25 *C. enderleini* (2 male), 10 *C. brucei* and one each of *C. grahamii, C. miombo* and *C. kibatiensis*; all female unless specified), while the second largest catch was 107 at SH2 (87 *C. imicola* (3 male), 10 *C. bolitinos*, 5 *C. nivosus* (1 male), 3 *C. enderleini* (3 male) and one each of *C. brucei* and *C. similis*). In the facility itself the single largest nightly catch was 23 *Culicoides* (17 *C. similis*, 2 *C. imicola*, 3 *C. trifasciellus* (1 male) and 1 *C. neavei*) which were trapped near a pond behind one of the greenhouses, followed by 19 *Culicoides* (6 *C. trifasciellus*, 5 *C. bedfordi* (1 male), 3 *C. imicola*, 4 *C. enderleini* (1 male) and 1 *C. similis*) trapped at a leaf litter composting site between two of greenhouses. Inside the greenhouses themselves, we caught 9 and 2 *Culicoides* in two of the eight greenhouses. These comprised 7 *C. similis*, 1 *C. imicola*, 1 *C. pycnostictus* (all female) and one male each of *C. enderleini* and *C. bedfordi*. No *Culicoides* were caught in the remaining sites. The most abundant species of *Culicoides* were *C. imicola* (55.4 %), followed by *C. bolitinos* (24.1%), *C. enderleini* (6.6%) and *C. similis* (6.1%) (Table 1).

**Table 1.** *Culicoides* species trapped at a flower growing and packaging plant, and two adjacent smallholdings, in Kenya.

| Species Trapped | Female (% of total catch) | Male (% of total catch) |
| --- | --- | --- |
| *C. bedfordi* | 5 (1.2) | 2 (9.5) |
| *C. bolitinos* | 102 (24.1) | 0 |
| *C. brucei* | 11 (2.6) | 0 |
| *C. enderleini* | 28 (6.6) | 9 (42.9) |
| *C. grahamii* | 1 (0.2) | 0 |
| *C. imicola* | 235 (55.4) | 7 (33.3) |
| *C. kabatiensis* | 1 (0.2) | 0 |
| *C. miombo* | 1 (0.2) | 0 |
| *C. neavei* | 1 (0.2) | 0 |
| *C. nivosus* | 4 (0.9) | 1 (4.8) |
| *C. pycnostictus* | 1 (0.2) | 1 (4.8) |
| *C. similis* | 26 (6.1) | 0 |
| *C. trifasciellus* | 8 (1.9) | 1 (4.8) |
| **Total** | **424 (100)** | **21 (100)** |

Blood-fed members of *C. imicola* and *C bolitinos* were trapped solely on the smallholdings. All *C. imicola* trapped on the flower farm were gravid. No *C. bolitinos* were trapped in the flower farm.

Of the other insects trapped, the single largest catch (*n=*857) was also at SH2, with the second largest catch from near a pond behind the greenhouses (*n=*530). Non-*Culicoides* were trapped at every sampling location (including inside every greenhouse). The largest single catch inside a greenhouse was 97 non-*Culicoides*. This was the same trapping sample that produced the single largest catch of *Culicoides*. No *Culicoides* were trapped from aspirating directly from plants, or from the emergence traps.

Members of the Family Psychodidae were the most abundant non-*Culicoides* trapped (*n=*683; 26.10% of other insects), followed by the Family Chironomidae (*n=*515; 19.68%) (Table 2).

**Table 2.**
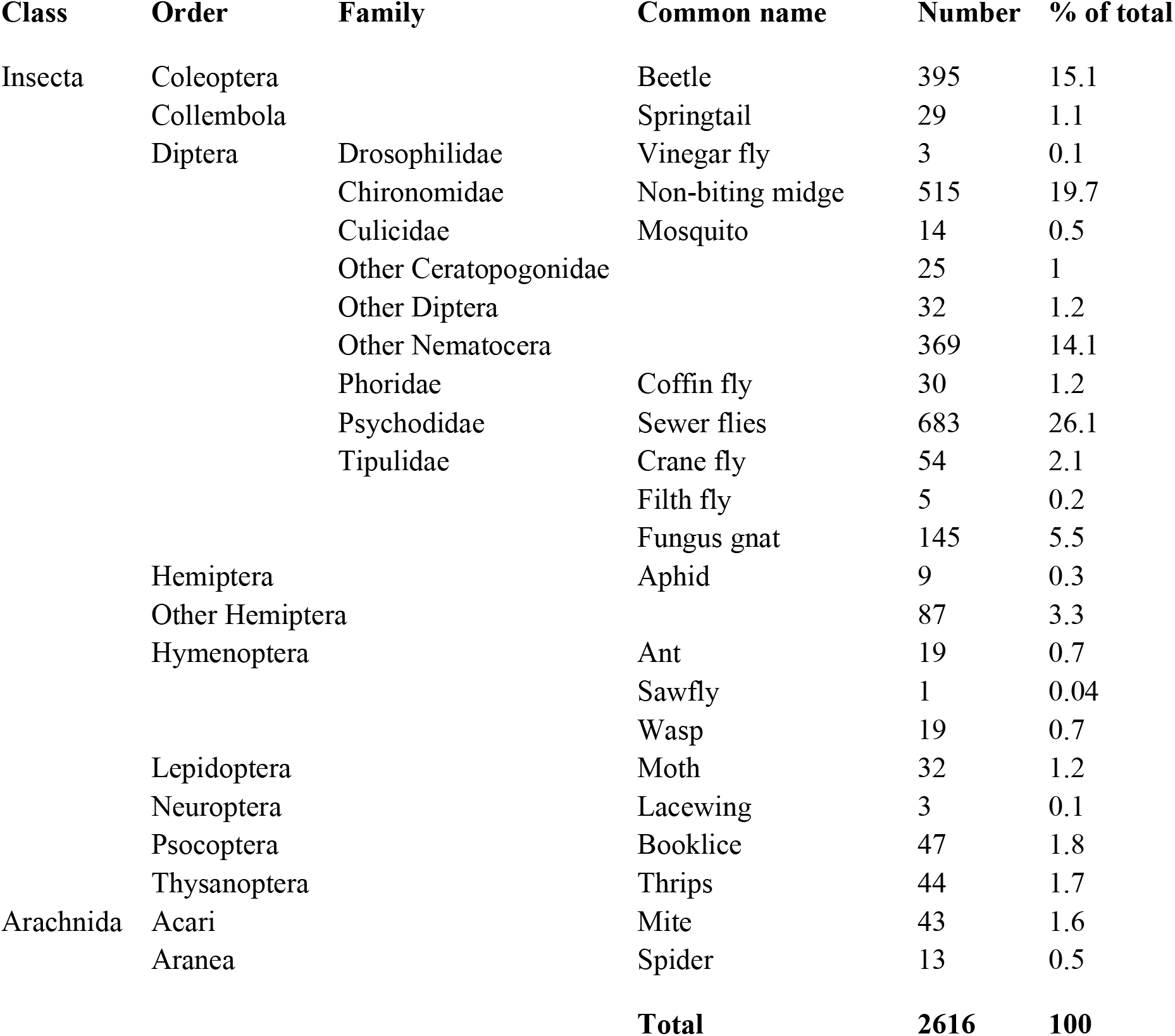
Other arthropods trapped at a flower growing and packaging plant, and two adjacent smallholdings, in Kenya.

### Stages of Production

Traps set outdoors captured the greatest numbers of *Culicoides* (SH1, *n=*269; SH2 *n=*122; near water sources, *n=*23; composting sites, *n=*19; rootstock, *n=*1). Traps set in greenhouses collected a total of only 11 *Culicoides*, while traps set in the packaging room, cold storage and transportation captured no *Culicoides* at all. Similar results were seen for the non-*Culicoides* trapped, with largest numbers captured in outdoor environments, and fewer captured as the stages of production neared cold storage and transportation (Table 3).

**Table 3.** *Culicoides* and other insects trapped at each stage of production on a flower growing and packaging plant in Kenya.

| Production Stage | Number of *Culicoides*<br>(% of total catch) | Number of Non-*Culicoides*<br>(% of total catch) |
| --- | --- | --- |
| Smallholding 1 | 269 (60.5) | 918 (35.1) |
| Smallholding 2 | 122 (27.4) | 454 (17.4) |
| Near Water Sources | 23 (5.2) | 530 (20.3) |
| Composting | 19 (4.3) | 185 (7.1) |
| Rootstock/ outdoor crops | 1 (0.2) | 90 (3.4) |
| Indoor Propagation* | 0 | 91 (3.5) |
| Growing on* | 11 (2.5) | 177 (6.8) |
| Packaging Room | 0 | 166 (6.3) |
| Cold Storage Room | 0 | 0 |
| Transportation | 0 | 6 (0.2) |
| **Total** | **445 (100)** | **2617 (100)** |

### Disease and Pest Control

Both mechanical and chemical pest control methods were undertaken at the site. Three of the greenhouses sampled had mechanical pest control options in place. These included roller traps for white flies, aphids, leaf miners and thrips; moth and butterfly traps, fruit fly traps, pest level monitoring traps and scented thrips traps (Figure 1). Air circulation systems were in place and insecticides sprayed overnight in greenhouses depending on what pests were found to be present. A number of the greenhouses were also used to test biological control methods for pests. Of the greenhouses sampled, three of them were sprayed with insecticide the night before trapping took place; with 3 greenhouses recording details of known pests and diseases present (Table 4).

**Table 4.**
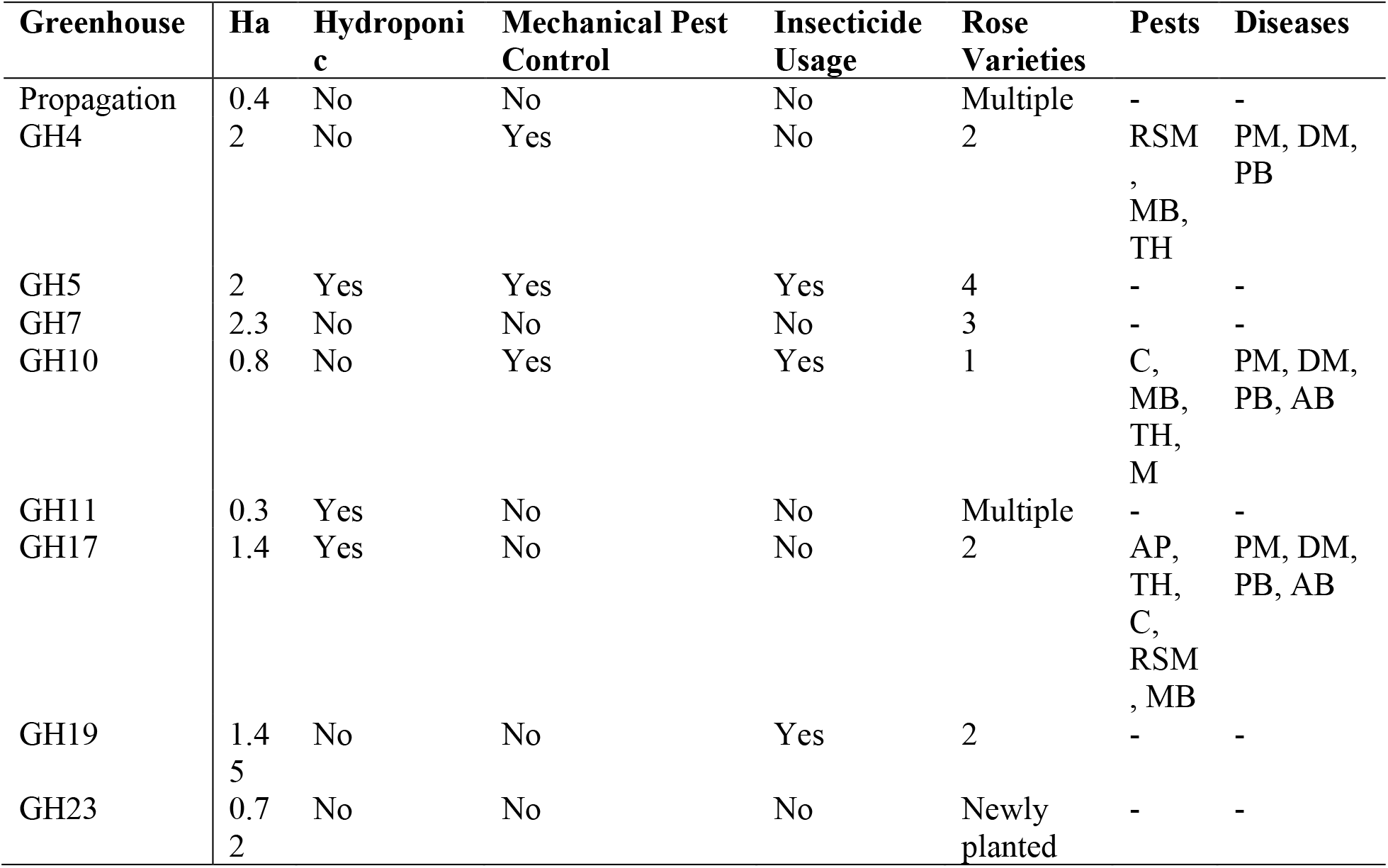
The eight growing greenhouses and one propagation greenhouse sampled at a flower farm in Kenya. Details given on: the number of hectares occupied by the greenhouse (Ha); whether hydroponics or mechanical pest management methods were employed; whether the greenhouse was sprayed with insecticide prior to sampling; the pests (RSM = red spider mite; MB = mealybugs; TH = thrips; C = caterpillars; M = mites; AP = aphids) and diseases (PM = powdery mildew; DM = downy mildew; PB = petal botrytis; AB = agrobacteria) to be controlled.

**Figure 1.**
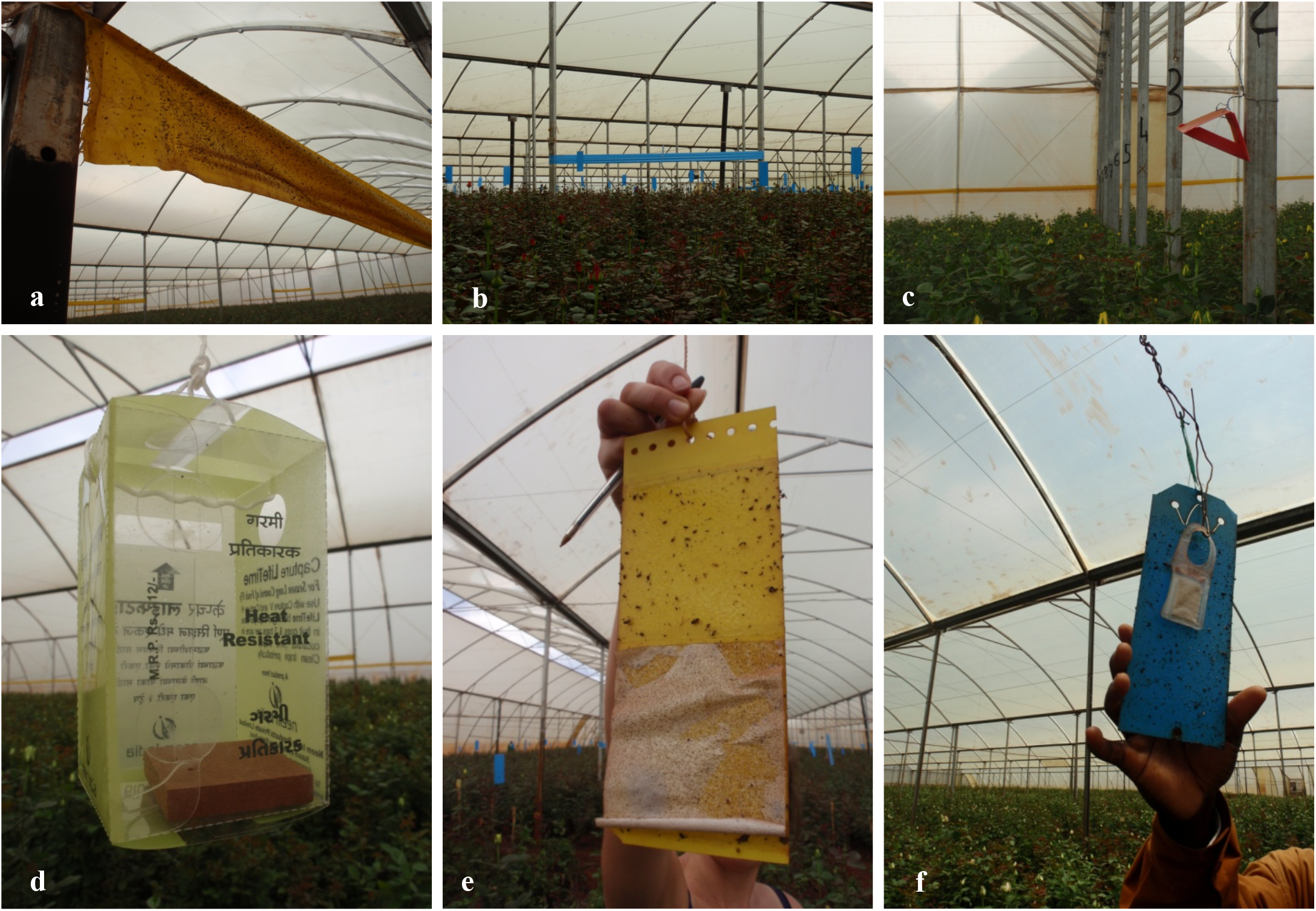
Methods of mechanical pest control undertaken within the greenhouses at a flower-packaging plant in Kenya. a. roller trap for white flies, aphids, leaf miners; b. roller trap for thrips; c. moth & butterfly traps; d. fruit fly traps; e. pest-level monitoring traps; and f. scented thrips traps.

## Discussion

This study is the first to investigate whether *Culicoides* biting midges are present on flower farms and packaging plants in Africa. The results demonstrate that *Culicoides* could be found in substantial numbers on smallholdings adjacent to the flower farm, and that small numbers of *Culicoides* were caught in the greenhouses themselves. However, none were trapped where cut flowers were packaged for export. The species caught in the smallholdings and greenhouses included the major Afrotropical vector of several arboviruses, *C. imicola*; while the established vector of bluetongue virus and African horse sickness virus, *C. bolitinos*, was trapped in the smallholdings but not in the greenhouses. We failed to trap any *Culicoides* at all in the areas where cut flowers were prepared for export. However, given the scale of the flower trade from Africa to Europe, our results are consistent with there being a very small, but non-zero risk of biting midges occasionally being transported along with the produce.

### Route of Introduction to Europe

There are several hypotheses as to the possible introduction of *Culicoides* into northern Europe in 2006. One theory is that infected *Culicoides* were transported into the Maastricht area of the Netherlands with cut flowers from Africa. The insects are hypothesised to have either entered consignments during packaging or shipping, or to have been phytotelmatic species which breed exclusively in container-habitats, including water in the leaf axils of plants. Only 5% of *Culicoides* species are phytotelmatic however, and of those species only C. *loughnani* is said to have a substantiated record of long-distance introduction on plants [15]. This contrasts with other insects such as the mosquito *Aedes albopictus*, where their worldwide transportation, in lucky bamboo plants imported from south-east Asia, is well documented. Even if there is phytotelmatic transport of juvenile *Culicoides*, the appearance of BTV-8 in northern Europe remains enigmatic as, in the absence of documented vertical transmission of BTV, only adult female *Culicoides* can be infected with such viruses.

Kenya is the third largest flower exporter globally, after Colombia and Ecuador, and is the lead exporter of cut roses to the EU (38%). Of these flower consignments, almost 65% are sold through Dutch auction houses, with the industry seeing a growth of 5^%^ each year. The EU absorbs 85% of all flower exports from Kenya, with its main cut flower export being roses (74% of exports), and followed by *Dianthus caryophyllus* (carnations) and *Alstroemeria* (Peruvian lily). A Kenyan rose-producer and exporter is therefore the ideal field site for assessing whether female *Culicoides* can be found within flower-packaging plants, as well as the level of risk of *Culicoides* being transported in cut flowers to Europe.

### Environment at a Flower Farm

An important aspect of the current study is that midges were trapped at different stages of the production process, which highlights that although *Culicoides* are present around the flower farm, their numbers diminish with each stage of the production process, therefore reducing the likelihood of transporting them with each stage of the process.

According to [11], consignments of plants are usually packed in the country of origin at night, when conditions (lighting, humidity) during packaging would appear favourable for incorporating insects during packaging, and plants are watered prior to shipment. This was not the case for the flower farm in Kenya where this trapping took place. During daytime, cut flowers were held in water until packaged, and were then stored in a sealed room at 4°C until transportation to the airport. The flowers were transported in refrigerated lorries directly to the airport prior to shipment. No flowers were processed or transported at night. We do not know if this is now standard practice in Kenya but, in our view, this approach presents a lower risk of transporting *Culicoides* than preparing cut flowers at night.

### *Culicoides* Trapped

Of the 75 arboviruses associated with *Culicoides*, 15 have been isolated from species belonging to *C. imicola* complex [1]. The *C. imicola* complex (*C. imicola, C. bolitinos, C. brevitarsis, C. nudipalpis, C. kwagga, C. loxodontis, C. miombo, C*.*pseudopallidipennis, C. tutti-frutti* and *C. asiana*) [2, 3] contains the three most important known vectors of bluetongue virus: *C. imicola, C. bolitinos* and *C. brevitarsis*. Both *C. imicola* and *C. bolitinos* were trapped during this study. *C. bolitinos* was trapped exclusively on both smallholdings outside the flower farm, likely due to its strong association with cattle dung as a breeding site. *C. imicola* was trapped at both smallholdings, but also trapped near to a pond, at a composting site, and inside two greenhouses on the flower farm. This may be an indication of their need for semi-moist soil breeding sites, as opposed to the dung sites *C. bolitinos* occupy.

## Conclusions

While we did not detect any *Culicoides* within the facility where cut flowers are prepared for export, we did detect low numbers of several species in the greenhouses where the flowers are grown, including some individuals of an important arbovirus vector. Significant numbers of *Culicoides*, including two vector species, were detected in close vicinity to the farm where livestock were kept. While not directly evidenced, our data are consistent with a small but non-zero risk of *Culicoides* vectors occasionally being transported along with cut flowers. Given the scale and value of the international industry in flower exports, and the potential risks to importing countries, we recommend simple interventions that reduce risks and protect the industry. This should include light traps for biting midges in the processing rooms of flower farms and importing airports. We further recommend that flower farms work with their smallholder neighbours to remove *Culicoides* breeding sites and/or hang midge-proof netting (with a minimum 1000 holes per square inch) on the side of greenhouses close to smallholdings.

## Author Contributions

The study was conceived and designed by JS, EF and MB. *Culicoides* were collected by JS in Kenya. EF selected the sampling location and provided consumables while in Kenya. Species-level identifications were undertaken by KL. All authors contributed to the final version of the manuscript.

## Acknowledgements

This study is funded by University of Liverpool Living with Environmental Change (LWEC) pump priming, carried out by GK & JS under the supervision of MB. JS was a BBSRC-funded DTP student. This work was undertaken as part of a collaborative project with EF (ILRI) and KL (ARC). The authors would like to acknowledge the flower farm where sampling was undertaken and Dr Georgette Kluiters for the significant contribution she made to the study’s design and execution.

